# DIFFERENTIAL ACTIVATION OF G-PROTEIN SUBTYPES BY DOPAMINE D1 RECEPTOR ALLOSTERIC MODULATION

**DOI:** 10.64898/2026.09.26.754724

**Authors:** Anh Minh Nguyen, Hayato Umaoka, Hideaki Yano

## Abstract

Allosteric modulation of G protein-coupled receptors (GPCR) is a rapidly advancing area of drug discovery with considerable therapeutic potential. In this study, we used various bioluminescence resonance energy transfer (BRET) assay configurations to investigate the differential activity of LY3154207, a dopamine D1 receptor (D1R)-positive allosteric modulator (PAM), in potentiating D1R-mediated signaling pathways. Our data showed that LY3154207 demonstrated clear bias toward β-arrestin recruitment over G-protein engagement. Among the G-protein subtypes, LY3154207 preferentially potentiated Gα_s_ signaling induced by DA. Interestingly, when tested for its effect on the Gα_s_-selective D1R agonist DHX and the Gα_olf_-selective agonist TAV, LY3154207 further enhanced the selectivity of these compounds, suggesting a probe-dependent effect. We also studied the structural mechanism underlying this subtype selectivity and found that W123 and R130 in the intracellular loop 2 (ICL2) of D1R were crucial for the activity of LY3154207. Furthermore, the αN domain of the G-protein appeared to make minor contributions to this G-protein subtype selectivity of LY3154207. Lastly, our data demonstrated that LY3154207 induced cAMP production in the absence of the orthosteric ligand, indicating that this compound is an ago-PAM.

## 1. INTRODUCTION

Dopamine D1 receptor (D1R) is the most abundant DA receptor in the CNS and mediates the outputs of most dopaminergic pathways.^1,2^ Its involvement in controlling crucial behaviors, including voluntary movement, substance abuse, and other cognitive functions, makes D1R a promising target for drug development against several neurological and neuropsychiatric diseases.^3–8^ However, developing selective D1R agonists has proven to be challenging. Most D1R agonists displayed unfavorable pharmacokinetics, tolerance, and detrimental adverse effects, such as epileptogenicity and hypotension.^9–14^

Selectively activating D1R in the prefrontal cortex (PFC) or the striatum has been suggested to be beneficial in treating the negative symptoms of schizophrenia (SZ) or Parkinson’s disease (PD), respectively.^15^ We posited that region-specific pharmacology in D1R could be achieved by taking advantage of the contrast localization of the two G-proteins coupling with D1R, where Gα_s_ is highly expressed in the cortical regions and Gα_olf_ is predominantly found in the striatal areas.^16^ Our previous publication supported this hypothesis and demonstrated that D1R agonists that induced preferential engagement of one G-protein could selectively modulate the electrophysiological activities and behaviors associated with the brain region where that G-protein was distributed.^17,18^

Positive allosteric modulators (PAM) represent a new frontier in drug development. These compounds bind to the receptor in pockets spatially distinct from the orthosteric binding site, enhancing the endogenous ligand’s affinity for the receptor and thereby increasing the cellular signaling induced by the orthosteric ligand.^19^ Importantly, PAMs preserve the spatial and temporal aspects of the endogenous ligands, allowing select signaling pathways to be activated in their physiological pattern.^20^ Notably, the allosteric binding pockets are usually less conserved than the orthosteric pocket; thus, targeting these sites might confer better receptor subtype selectivity.^21^ In addition, one receptor can have more than one allosteric binding site, and these sites can be occupied simultaneously to synergistically affect the receptor’s affinity toward its endogenous ligand.^22–24^ Of these allosteric sites, the pocket formed by transmembrane domain 3 (TM3), intracellular loop 2 (ICL2), and TM4 (hereafter referred to as the ICL2 pocket) has been widely studied. Although this is an allosteric pocket shared by several GPCRs, including D1R and β2-adrenergic receptor (β2AR),^25,26^ its sequence is highly diverse amongst these receptors, which facilitates developing receptor-selective drugs.^27^ In addition, targeting the ICL2 pocket can stabilize the receptor in specific conformations that favor different transducers, thereby inducing signaling bias.^28,29^

LY3154207 is a PAM that binds to D1R at the ICL2 pocket and potentiates both G-protein and β-arrestin signaling induced by D1R agonists.^27^ This compound promotes wakefulness in humanized D1R mice and sleep-deprived male volunteers.^30^ Data from phase I (NCT03616795, NCT02562768) and phase II (NCT03305809) trials reported that this compound was well tolerated by both healthy subjects and patients and that it significantly improved clinical scores in patients with Alzheimer’s disease.^31–33^ These reports emphasized the potential of PAMs as an alternative to traditional dopaminergic therapy. Built on our previous report, we investigated whether LY3154207 showed preference in potentiating the signaling cascade mediated by one G-protein over the other. We employed bioluminescence resonant energy transfer (BRET) assays to study the potentiating effect of LY3154207 on cAMP production induced by dopamine (DA), the Gα_s_-biased agonist dihydrexidine (DHX), and the Gα_olf_-biased agonist tavapadon (TAV).^17,18^ We then used site-directed mutagenesis followed by functional BRET assays to reveal the residues in LY3154207 binding pocket crucial to its allosteric modulating effect and to elucidate the molecular interactions at the receptor-G-protein interface governing this subtype selectivity. Our results demonstrated that G-protein subtype selectivity can be exploited in D1R PAMs, opening a new trajectory in drug development against challenging neurological and neuropsychiatric diseases.

## 2. MATERIALS AND METHODS

### 2.1. Materials

*Reagents for cell culture:* Dulbecco’s modified Eagle’s medium (DMEM), fetal bovine serum (FBS), L-glutamine, penicillin-streptomycin

*Reagents for transfection*: polyethyleneimine (PEI)

*Reagents for BRET*: KCl, NaCl, MgCl_2_, K-EGTA, Na-HEPES, NaHSO3, phosphate-buffered saline (PBS), h-coelenterazine

*Drugs:* DA, DHX, TAV, and LY3154207. Drugs were prepared in 10 mM stock in DMSO and stored at −20 °C until use.

### 2.2. Site-directed Mutagenesis

Site-directed mutagenesis was performed following the instructions from the QuikChange Lightning Site-Directed Mutagenesis Kit by Agilent. In short, a 25 μl reaction was set up with a 1:1:1 ratio between the plasmid template and the two primers, which were previously designed and synthesized. The reaction was incubated in a thermal cycler for 18 cycles of 20 seconds at 95 °C, 10 seconds at 55 °C, and 3-4 minutes at 68 °C, followed by a 5-minute incubation period at 37 °C with *Dpn* I to digest the original template.

The mixture was transformed into Stellar competent cells from Takara Bio by heat shock at 42 °C. After an hour of constant shaking at 230 rpm at 37 °C to recover, the cells were plated on Luria-Bertani (LB) agar supplemented with 100 μg/ml ampicillin. At least four colonies on the plate were collected and extracted for their plasmid using the QIAprep Spin Miniprep Kit from QIAGEN. Successfully mutated plasmids were confirmed by sequencing.

### 2.3. Cell Culture

Gα_s/olf_-KO HEK293 cells were cultured in DMEM containing 10% FBS, 2 mM L-glutamine, and 1% penicillin-streptomycin at 37 °C and incubated in 5% CO_2_ −95% moisturized air.

### 2.4. Transfection

Cells were seeded at 4 million cells per 10-cm plate. A constant amount of cDNA was transfected using PEI (1:2 w/w). The ratio between the transfected constructs had been previously optimized for the dynamic range of the drug-induced BRET change, as described previously.^18^ All cDNAs were prepared using the QIAGEN Plasmid Maxi/Midi kit.

*For engagement BRET assays*, cells were transfected with D1R-RLuc8 (Renilla Luciferase), Gα_s_-154Venus, Gβ2, Gγ7, and RIC8B.

*For β-arrestin recruitment assays*, cells were transfected with D1R-RLuc8, β-arrestin2-Venus, and GRK2.

*For cAMP production assays*, cells were transfected with D1R or D1R mutants, WT or chimeric Gα_s_ or Gα_olf_, Gβ2, Gγ7, RIC8B, and the biosensor CAMYEL.

### 2.5. Bioluminescence Resonant Energy Transfer (BRET) Assay

Forty-eight hours post-transfection, cells were harvested and resuspended in potassium-based buffer (KPS) (140 mM KCl, 10 mM NaCl, 1 mM MgCl_2_, 0.1 mM K-EGTA, 20 mM Na-HEPES, pH 7.2) for engagement BRET or phosphate-buffered saline (PBS) containing 200 μM sodium bisulfite (NaHSO_3_) for β-arrestin recruitment assays and cAMP assay. Cells were distributed at 500000 cells per well in a white 96-well plate.

Cells were incubated with LY3154207 at varying concentrations for 30 mins. Then, 5 μM coelenterazine H was added. D1R agonists at varying concentrations were added 1 minute later. Bioluminescence and fluorescence emissions were recorded at 485 and 530 nm, respectively.

### 2.6. Bias Factor Calculation

Bias factor between Gα_s_ and β-arrestin is calculated as previously described using the following equation.^34,35^

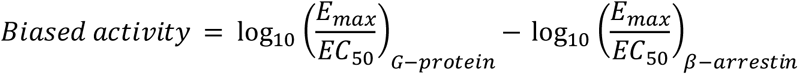

### 2.7. Data and Statistical Analysis

All experiments were conducted at least 5 times independently. In every experiment, each concentration constituting the dose-response curve was run in triplicate. The value of n represents the number of independent experiments. Data were normalized to the E_max_ of the D1R agonists in the absence of LY3154207 in that condition and presented as mean ± S.E.M (standard error of the mean). One-way ANOVA followed by Dunnett post-hoc analysis was employed to determine statistical significance (P<0.05). Data presentation and statistical analysis were performed using GraphPad Prism 10.

## 3. RESULTS

### 3.1. LY3154207 potentiated different D1R-mediated signaling pathways

LY3154207 has been reported to potentiate various D1R signaling pathways.^27^ Using bioluminescence resonance energy transfer (BRET) assay, LY3154207 was evaluated for its effect on enhancing the coupling of D1R with two of its major transducers, Gα_s_ and β-arrestin. Specifically, D1R harbored Renilla luciferase, while both transducers carried the yellow fluorescent protein, Venus. The positions where the donor and the acceptor of the BRET pair were tagged onto the receptor and the transducers were experimentally determined.^36^

LY3154207 potentiated the DA-induced engagement of Gα_s_ and recruitment of β-arrestin to D1R in a concentration-dependent manner **(Figure 1)**. In the Gα_s_ engagement assay, cells treated with LY3154207 at 1.0 µM showed a significant leftward shift of the DA dose-response curve (0.7-fold shift), and the maximal shift was observed at 100.0 µM LY3154207 (0.9-fold shift; **Figure 1A, Supplementary Table 1**). In addition, LY3154207 at 10.0 µM significantly increased the E_max_ of the DA dose-response curve (7.5% increase compared to DA alone; **Figure 1A, Supplementary Table 1**).

**Figure 1.**
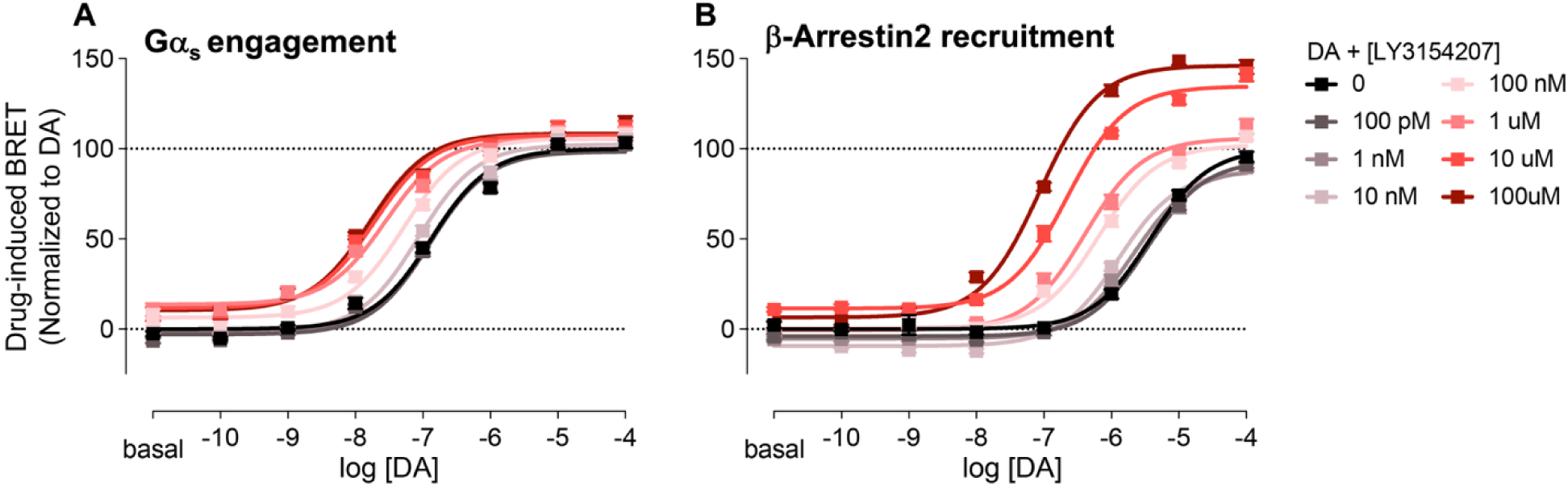
LY3154207 potentiated different signaling pathways in D1R. Drug-induced BRET ratio change between D1R-Rluc8 and Gα_s_-Venus **(A)** or β-arrestin-Venus **(B)**. Data were normalized to the E_max_ of DA alone in the same condition and presented as mean ± S.E.M (n ≥ 3)

Interestingly, LY3154207 appeared more potent at potentiating β-arrestin recruitment in D1R. The compound significantly shifted the DA dose-response curve leftward at 100.0 nM (0.8-fold shift). This effect increased proportionally as the concentration of LY3154207 was raised. At 100.0 µM, LY3154207 caused a dramatic 1.6-fold leftward shift of the DA dose-response curve **(Figure 1B, Supplementary Table 1)**. Consistent with its effect on DA potency, LY3154207 significantly potentiated DA efficacy, increasing the E_max_ of DA by 35% and 46% when cells were pretreated with 10.0 and 100.0 µM LY3154207, respectively **(Figure 1B, Supplementary Table 1)**. Calculating the bias factor between both assays at 100.0 µM LY3154207 suggested that LY3154207 was biased toward β-arrestin (ΔΔlog = −0.86**; Supplementary Table 2**).

### 3.2. LY3154207 differentially modulated DA-induced D1R-mediated signaling through Gα_s_ and Gα _olf_

We previously reported that G-protein subtype selectivity in D1R agonists might be exploited to selectively modulate physiological functions associated with the brain region where Gα_s_ and Gα_olf_ were distributed.^17^ Here, we asked whether LY3154207’s success in clinical trials might be partly attributed to its preference for promoting coupling of D1R toward one specific G-protein subtype. Thus, we evaluated the potentiating effect of LY3154207 on DA-induced G-protein engagement and cAMP production mediated through both subtypes.

To maintain consistency in the G-protein constructs used for both assays, we designed an engagement BRET configuration in which luciferase was tagged on γ7 subunit of the G-protein trimer and Venus was carried by D1R. In both G-proteins, LY3154207 induced a left shift of the DA dose-response curve in a concentration-related fashion **(Figure 2A, 2B)**. LY3154207 significantly shifted the DA dose-response curve leftward in Gα_s_ (0.76-fold) and Gα_olf_ (0.70-fold) at 100.0 nM and reached the maximal potentiating activity for both G-proteins at 1.0 µM (0.80-fold in Gα_s_ and 0.94-fold in Gα_olf_; **Supplementary Table 3**). Contrary to its effect on potency, LY3154207 did not alter the E_max_ of DA in the Gα_olf_ engagement assay at any concentrations tested **(Figure 2B)**. However, it induced a 20% increase in the E_max_ of DA in promoting Gα_s_ coupling, although this change was not statistically significant **(Figure 2A)**.

**Figure 2.**
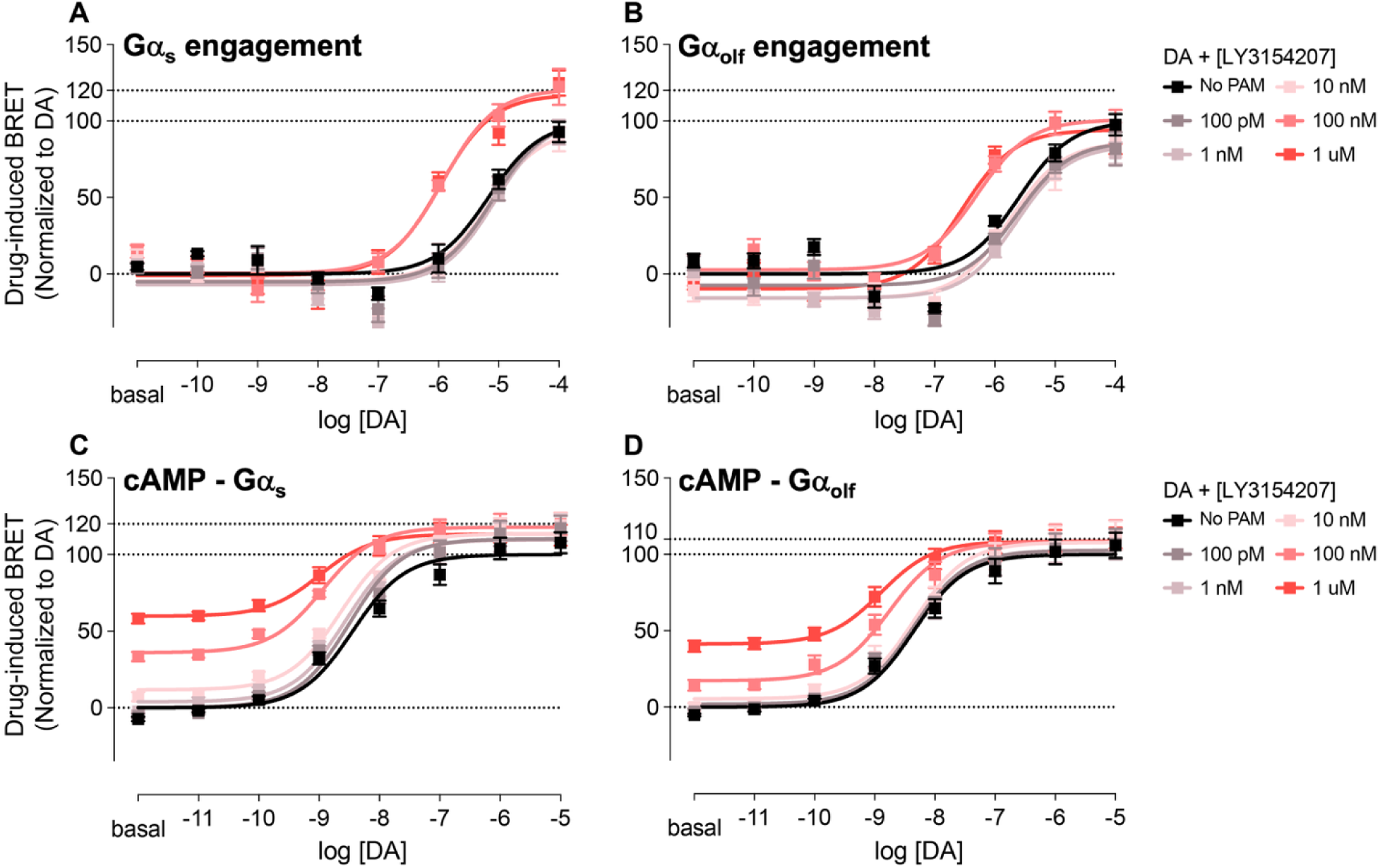
**LY3154207 differentially potentiated D1R signaling mediated by Gα_s_ and Gα _olf_** Drug-induced BRET ratio change between D1R-Venus and Gα_s_-γ7-Rluc8 **(A)** or Gα_olf_-γ7-Rluc8 **(B)**. Drug-induced BRET ratio change of the cAMP biosensor CAMYEL mediated by Gα_s_ **(C)** or Gα_olf_ **(D)**. Data were normalized to the E_max_ of DA alone in the same condition and presented as mean ± S.E.M (n ≥ 3)

To monitor the intracellular cAMP level, we used the BRET-based cAMP biosensor CAMYEL. LY3154207 dose-dependently potentiated the potency of DA-induced cAMP production via both G-proteins, with a maximal leftward shift of the DA dose-response curve achieved at 1.0 µM (0.55-fold in both Gα_s_ and Gα_olf_; **Figure 2C, 2D, Supplementary Table 4**). In addition, there was a clear separation between the DA dose-response curves separated clearly in Gα_s_ at an LY3154207 concentration as low as 10.0 nM, whereas the leftward shift in Gα_olf_ occurred at 100.0 nM. This suggested a preference of LY3154207 for potentiating Gα_s_ signaling. Moreover, LY3154207 enhanced DA efficacy in a concentration-dependent manner in both G-proteins. Consistent with the engagement assay, LY3154207 showed a stronger effect in potentiating DA efficacy in Gα_s_, boosting a significant 18% increase in the E_max_ of DA in Gα_s_ at 100.0 nM compared to an 8% increase in Gα_olf_ **(Supplementary Table 4)**. This result suggested that LY3154207 preferentially potentiated D1R signaling via Gα_s_ over Gα_olf_. Interestingly, LY3154207 can induce cAMP production in a dose-dependent manner in both G-proteins in the absence of the orthosteric ligand, with higher efficacy in Gα_s_ than in Gα_olf_, suggesting a preferential intrinsic activity of this compound toward Gα_s_ **(Figure 2C, 2D)**.

### 3.3. LY3154207 potentiated the effects of other D1R agonists

We previously reported that dihydrexidine (DHX) and tavapadon (TAV) induced preferential coupling of D1R to Gα_s_ and Gα_olf_, respectively.^17^ Given its selective effect on endogenous ligand activity **(Figure 2)**, we investigated how LY3154207 would act on these D1R agonists with distinct pharmacological profiles using the cAMP assay. Consistent with the cAMP assay results with DA, LY3154207 dose-dependently elicited cAMP production without an orthosteric ligand **(Figure 3, 4)**. Importantly, LY3154207-induced intracellular cAMP levels were higher when mediated via Gα_s_. These results further supported that LY3154207 preferred signaling through Gα_s_.

**Figure 3.**
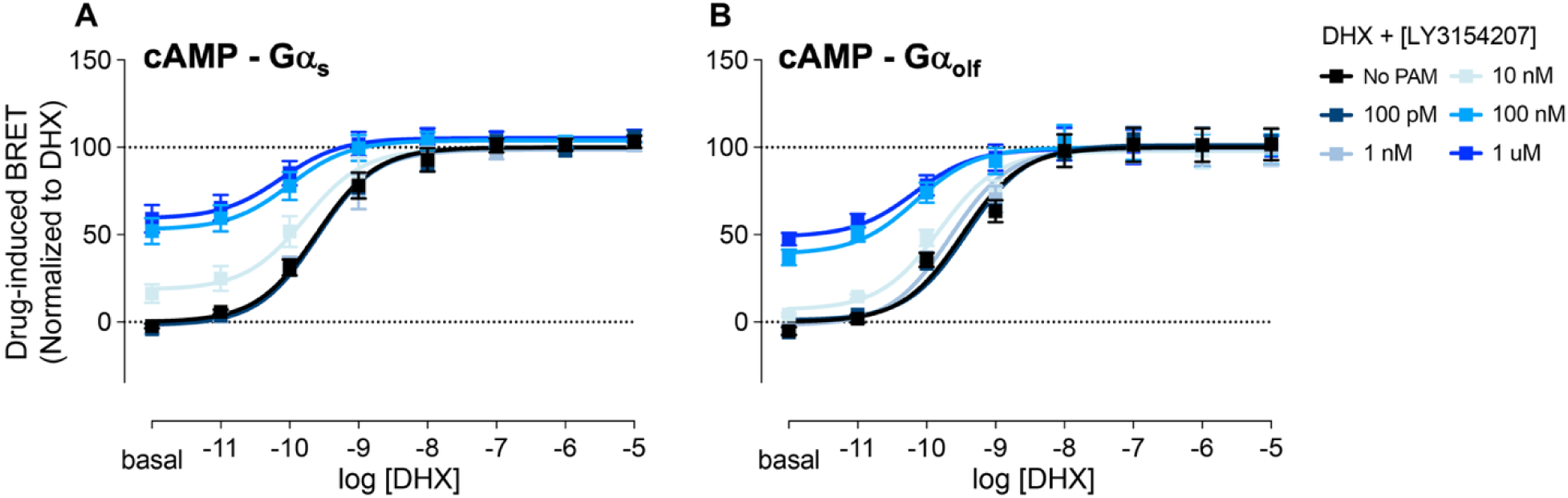
LY3154207 potentiated DHX-induced signaling in D1R. Drug-induced BRET ratio change of the cAMP biosensor CAMYEL mediated by Gα_s_ **(A)** or Gα_olf_ **(B)**. Data were normalized to the E_max_ of DHX alone in the same condition and presented as mean ± S.E.M (n ≥ 3)

**Figure 4.**
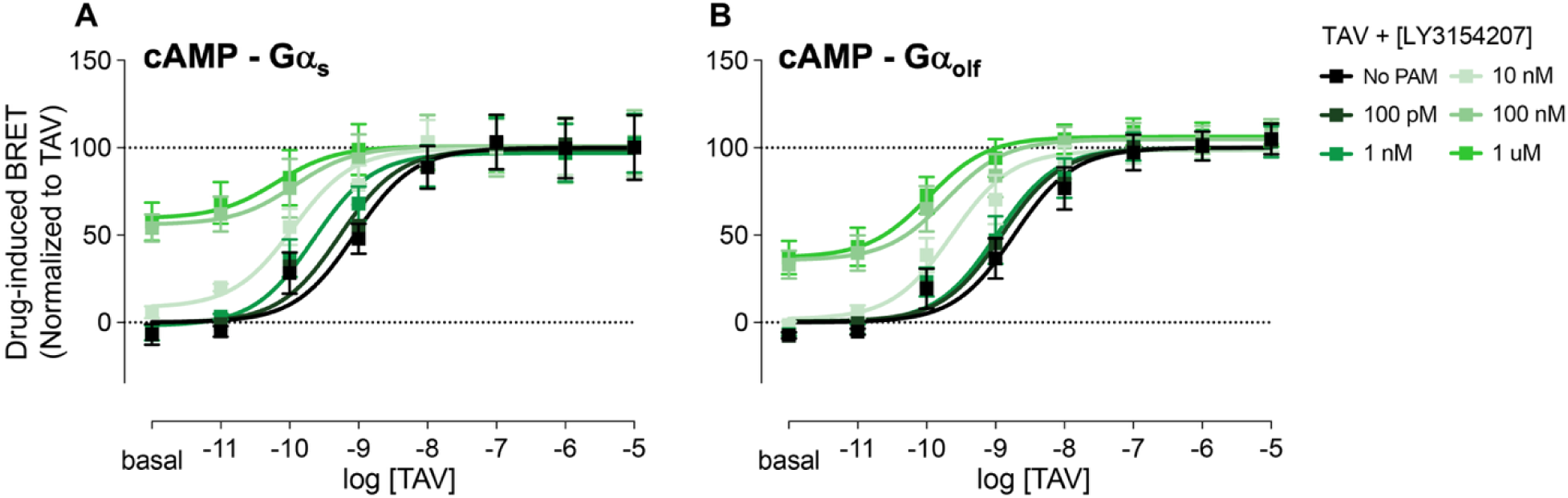
LY3154207 potentiated TAV-induced signaling in D1R. Drug-induced BRET ratio change of the cAMP biosensor CAMYEL mediated by Gα_s_ **(A)** or Gα_olf_ **(B)**. Data were normalized to the E_max_ of TAV alone in the same condition and presented as mean ± S.E.M (n ≥ 3)

LY3154207 potentiated DHX potency in both Gα_s_ and Gα_olf_ in a dose-dependent manner **(Figure 3A, 3B)**. The maximal potency shift was achieved at 1.0 µM in both G-proteins, albeit to different degrees (0.51-fold in Gα_s_ and 0.70-fold in Gα_olf_; **Supplementary Table 5**). In addition, the DHX dose-response curve in Gα_olf_ started shifting leftward at 1.0 nM LY3154207, while this shift in Gα_s_ did not occur until 10.0 nM, suggesting a higher affinity of LY3154207 for Gα_olf_ signaling induced by DHX. In contrast, LY3154207 1.0 µM increased DHX efficacy in Gα_s_ by 5% but failed to do so at every concentration tested in Gα_olf_ **(Figure 3A, 3B, Supplementary Table 5)**. These data implied that LY3154207 demonstrated a higher efficacy in potentiating DHX E_max_ in Gα_s_ over Gα_olf_.

LY3154207 dose-dependently shifted the TAV dose-response curves leftward in both G-proteins, with a maximal shift obtained at 1.0 µM (1.13-fold in Gα_s_ and 1.25-fold in Gα_olf_; **Figure 4A, 4B, Supplementary Table 6)**. In contrast to the results from the assays with DHX, a clear separation of the TAV dose-response curves in Gα_s_ was recorded at 1.0 nM LY3154207, while these events in Gα_olf_ occurred at an LY3154207 concentration of 10.0 nM, suggesting a higher affinity of LY3154207 for Gα_s_ signaling promoted by TAV. Interestingly, LY3154207 1.0 µM potentiated TAV efficacy in Gα_olf_ by 6% but not in Gα_s_, indicating a preferential efficacy-potentiating effect of LY3154207 for Gα_olf_ signaling triggered by TAV **(Supplementary Table 6)**. Taken together, these results revealed a unique characteristic of LY3154207 in enhancing the activity of D1R agonists with distinct G-protein subtype selectivity profiles.

### 3.4. Interaction with ICL2 was crucial for the effect of LY3154207

LY3154207 binds to D1R in a pocket formed by regions of TM3, TM4, TM5, and ICL2. In this pocket, LY3154207 assumes a horizontal pose, parallel to ICL2, with the dichlorophenyl ring sandwiched between an aromatic interaction with W123^3.52^ and a cation-π interaction with R130^ICL^^2^.^25,37,38^ In this study, we examined how W123A and R130A mutations at D1R affected LY3154207 potentiating activity on DA-induced signaling.

The W123A mutation did not interfere with DA activity at the mutant receptor. However, the potentiating effect of LY3154207 on the potency and efficacy of the DA-induced cAMP production was abrogated at all LY3154207 concentrations tested in both G-proteins **(Figure 5A, 5B, Supplementary Table 7, 8)**. In addition, W123A abolished LY3154207’s intrinsic activity to induce cAMP production in the absence of DA **(Figure 5A, 5B)**. This result suggested that an intact interaction with W123 was necessary for LY3154207 to exert its effect.

**Figure 5.**
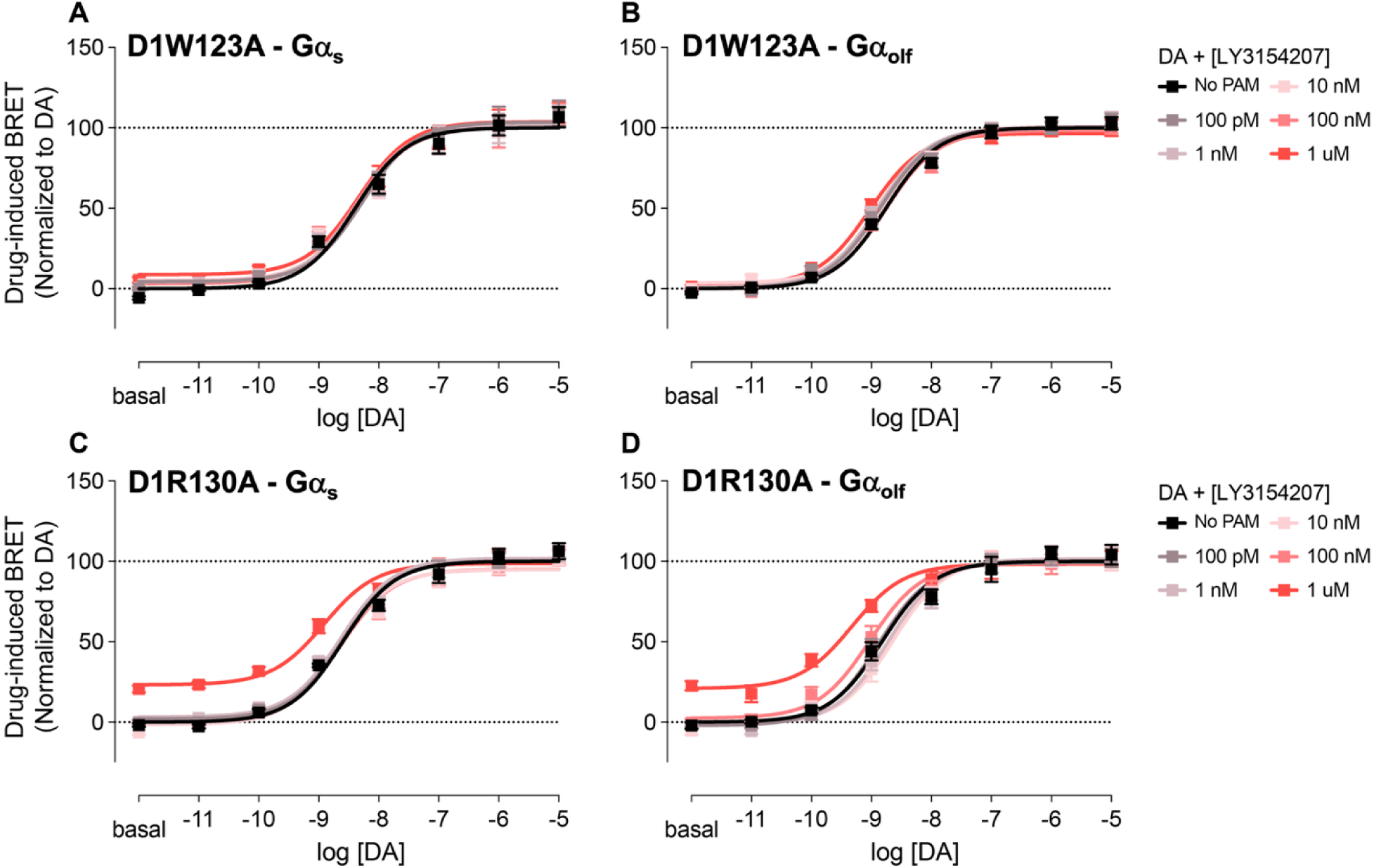
W123 and R130 were necessary for LY3154207 activity. Drug-induced BRET ratio change of the cAMP biosensor CAMYEL mediated by D1W123A-Gα_s_ **(A)**, D1W123A-Gα_olf_ **(B)**, D1R130A-Gα_s_ **(C)**, D1R130A-Gα_olf_ **(D)**. Data were normalized to the E_max_ of DA alone in the same condition and presented as mean ± S.E.M (n ≥ 3)

Similar to W123A, R130A did not affect the activity of the orthosteric ligand but drastically attenuated the potentiating effect of LY31 on DA in both G-proteins, albeit less severely than with W123A. None of the LY3154207 concentrations tested enhanced DA efficacy, and only at 1.0 µM did it induce a leftward shift in DA potency in both Gα_s_ and Gα_olf_, suggesting that R130 was important for LY3154207 activity **(Figure 5C, 5D, Supplementary Table 7, 8)**. Interestingly, this potency shift was less pronounced and not statistically significant compared to that reported for WT receptor with Gα_s_ (0.31-fold versus 0.55-fold, respectively) whereas in Gα_olf_, the leftward shift induced by 1.0 µM LY3154207 was comparable between the R130A and WT receptor **(Supplementary Table 4, 7, 8)**. This result implied that the Gα_s_-signaling pathway was more sensitive to changes in interactions between LY3154207 and the binding pocket.

### 3.5. The αN domain of G-protein contributed to the varying activity of LY3154207 at potentiating D1R-mediated signaling through Gα_s_ and Gα _olf_

αN plays a crucial role in G-protein recruitment, activation, and dissociation from the receptor.^39^ We previously reported that this domain contributed to the G-protein subtype selectivity of D1R induced by different D1R agonists.^17^ In this study, we investigated if this domain also affected the apparent preference of LY3154207 for Gα_s_ signaling. We generated two chimeric constructs of Gα_s_ and Gα_olf_ in which the six amino acids at the C-terminus of αN were swapped with their equivalent sequence on the other G-protein. We referred to the Gα_s_ carrying the six amino acids ERLAYK of Gα_olf_ αN as SOaN, and the Gα_olf_ whose αN contained the equivalent six amino acids DKQVYR from Gα_s_ as OSaN.

Both SOaN and OSaN retained their ability to mediate DA-induced cAMP production **(Figure 6A, 6B)**. LY3154207 potentiated the potency of the DA dose-response curves in both chimeric G-proteins in a concentration-dependent manner, to a level comparable to their WT counterparts **(Figure 6A, 6B, Supplementary Table 9)**. Furthermore, LY3154207 alone dose-dependently induced cAMP production via both chimeras, albeit with lower efficacy **(Figure 6A, 6B)**. However, LY3154207 did not show an efficacy-potentiating effect in either chimera, as all DA dose-response curves, with or without LY3154206, reached the same E_max_ **(Figure 6A, 6B)**. These data suggested a minor role of the αN domain in determining the apparent G-protein subtype selectivity seen with LY3154207.

**Figure 6.**
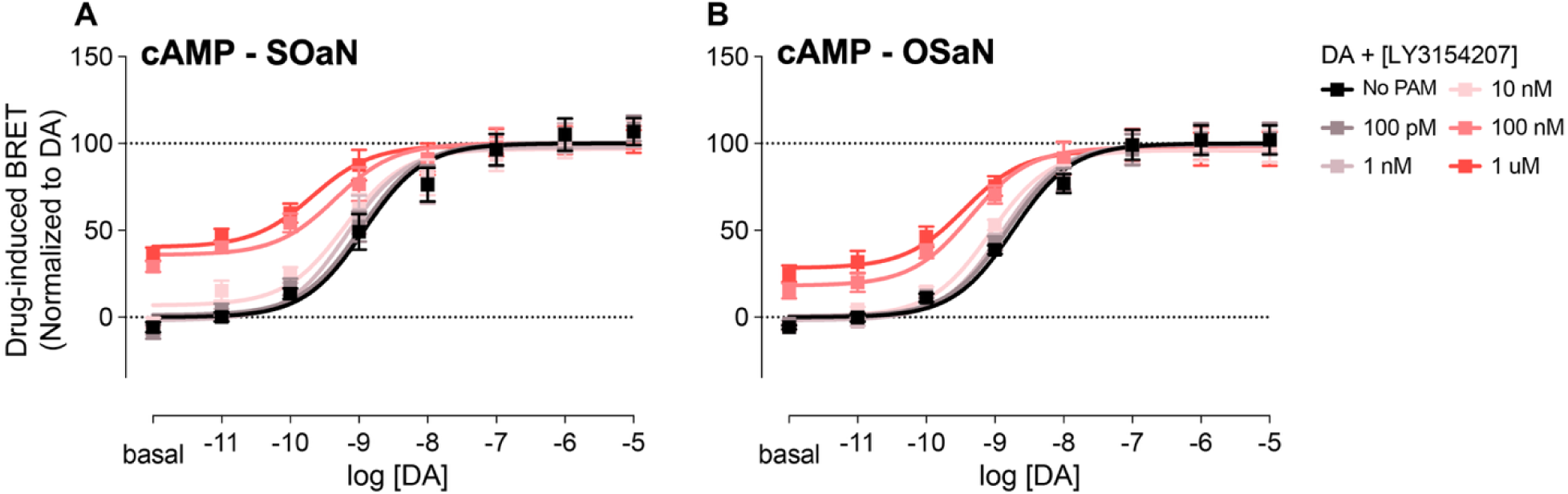
αN contributed to the subtype selectivity of LY3154207. Drug-induced BRET ratio change of the cAMP biosensor CAMYEL mediated by SOaN **(A)** or OSaN **(B)**. Data were normalized to the E_max_ of DA alone in the same condition and presented as mean ± S.E.M (n ≥ 3)

## 4. DISCUSSION

LY3154207 has been reported to potentiate various signaling pathways mediated by D1R.^27^ In this study, we used BRET to evaluate its effect on enhancing the DA-elicited coupling of D1R and two of its major transducers, Gα_s_ and β-arrestin, allowing us to directly investigate the interactions between the receptor and the transducers at a 1:1 ratio. LY3154207 induced leftward shifts in the DA dose-response curves in both pathways in a dose-dependent manner. Importantly, the shifts induced by LY3154207 at higher concentrations were considerably larger in β-arrestin recruitment than those in Gα_s_ engagement. Our bias factor calculation revealed a clear bias of LY3154207 for signaling through β-arrestin pathway, especially at higher concentrations of LY3154207. These data were in line with reports from other groups stating that LY3154207 was a β -arrestin-biased ligand.^27^ While this was beyond the scope of this study, LY3154207 could serve as a valuable lead compound for developing PAMs that selectively potentiate β-arrestin signaling.

While LY3154207 has been widely studied, to our knowledge, it has not been examined in the context of G protein subtype selectivity. The distinct distribution of Gα_s_ and Gα_olf_ in the brain opened up a potentially new approach to achieve region-specific pharmacology in D1R, which would be crucial in developing treatment for several detrimental neurological and neuropsychiatric disorders.^40–42^ Here, we evaluated LY3154207’s effect on the endogenous ligand DA’s ability to recruit G-proteins and promote cAMP production **(Figure 2)**. The lower potency of the DA curves in this engagement assay compared with **Figure 1** could be attributed to the different BRET pair used.^36^ Regardless, LY3154207 induced concentration-dependent, statistically significant leftward shifts in the DA dose-response curves for both G-proteins.

Although the potency shifts elicited by the compound were comparable between Gα_s_ and Gα_olf_, LY3154207 efficacy potentiation was only observed at Gα_s_, and not Gα_olf_, suggesting that the effect on efficacy was G-protein selective **(Figure 2A, 2B)**.

The cAMP assay agreed with our engagement BRET data. LY3154207 induced dose-dependent leftward shifts in the DA dose-response curves for both G-proteins with comparable effectiveness at the highest LY3154207 concentration tested **(Figure 2C, 2D)**. Interestingly, LY3154207 at lower concentrations appeared to be more effective at potentiating Gα_s_ signaling. In addition, while LY3154207 enhanced the E_max_ of DA in both G-proteins, it was more efficacious at Gα_s_. Taken together, these data indicated that LY3154207 preferentially potentiated Gα_s_ signaling elicited by DA.

In our previous publication, we showed that DHX was Gα_s_-selective and TAV was Gα_olf_-selective.^17^ Strikingly, LY3154207 demonstrated contrasting effects in potentiating signaling induced by these two compounds. With DHX, LY3154207 showed higher affinity for Gα_olf_ at low concentrations but selectively enhanced the E_max_ of DHX in Gα_s_ at the highest concentration tested **(Figure 3A, 3B)**. In contrast, the LY3154207-induced potency shift of TAV dose-response curves in Gα_s_ started at a lower concentration than in Gα_olf_, but TAV efficacy enhancement was observed only in Gα_olf_ **(Figure 4A, 4B)**. Taken together, these results suggested that the effect of LY3154207 on subtype-biased agonists was probe-dependent, and that it would intensify the bias level of the orthosteric ligand. To test this hypothesis, a larger sample size of D1R agonists with diverse bias profiles needs to be evaluated.

The aromatic interaction with W123^3.52^ and the cation-π interaction with R130^ICL^^2^ were necessary for LY3154207 to exert its effect.^25,37,38^ Using alanine scanning, we evaluated the cruciality of these residues to LY3154207 function. In both G-proteins, W123A did not affect the DA-induced cAMP production but completely abrogated the allosteric modulatory effect of LY3154206 to potentiate the potency and efficacy of the DA dose-response curves **(Figure 5)**. Furthermore, LY3154207 failed to induce cAMP production on its own in W123A, confirming the importance of this residue to the compound’s effect. Similarly, R130A drastically reduced LY3154207 activity as an ago-PAM without affecting the orthosteric ligand function. However, it retained some activity, although largely attenuated, at 1.0 µM **(Figure 5)**. Interestingly, the R130A mutation appeared to impact Gα_s_ signaling more severely than Gα_olf_, indicating that LY3154207 demonstrated a preferential effect toward Gα_s_ in DA-induced signaling. This provided valuable information for drug design, suggesting that modifying the strength of the interaction between the PAM and the residue would allow fine-tuning of the compound’s subtype selectivity. In addition to these two interactions, LY3154207 participated in extensive hydrophobic interactions in this pocket.^25,37,38^ Thus, investigating these residues would improve understanding of the mechanism by which LY3154207 exerts its G-protein subtype selectivity.

The αN domain of D1R participates in different stages of G-protein activation.^39^ We previously reported that this domain contributed to the subtype selectivity of D1R.^17^ By replacing the six amino acids at the C-terminus of αN in Gα_s_ and Gα_olf_ with the equivalent residues from the other G-protein, the agonism activities of DHX and TAV were significantly altered, trending toward the levels recorded in the WT G-protein from which the residues originated. In both SOaN and OSaN, LY3154207 lost its efficacy-potentiating effect on DA-induced cAMP but retained the ability to elicit leftward shifts in the DA dose-response curves compared to their WT G-proteins. Moreover, LY3154207-induced cAMP production was attenuated in both chimeric G-proteins. The mild changes in LY3154207 activity in SOaN and OSaN were not what we expected, given how significantly the efficacy of DHX and TAV were altered in these chimeras. These results suggest that other domains contribute to the G-protein subtype selectivity of LY3154207. Structural and molecular dynamics (MD) simulation studies are currently underway, which will provide further insights into the molecular mechanism of this phenomenon.

## 5. AUTHORSHIP CONTRIBUTION

**Anh Minh Nguyen**: Data curation, Formal analysis, Investigation, Visualization, Writing, **Hayato Umaoka**: Data curation, Formal analysis, Investigation, **Hideaki Yano**: Conceptualization, Data curation, Formal analysis, Funding acquisition, Resources, Supervision, Writing

## ACKNOWLEDGEMENTS

The authors would like to thank Dr. Asuka Inoue for kindly proving the HEK293 ΔGs cell line. This work is supported by Northeastern University Startup Funds (H.Y.).

## SUPPLEMENTARY DATA

**Supplementary Table 1.** LY3154207 potentiated D1R-mediated signaling pathways.

| DA + [LY31] | G$\alpha_s$ engagement | | β-arrestin2 recruitment | |
| --- | --- | --- | --- | --- |
|  | E<sub>max</sub> (%DA) | Log [EC<sub>50</sub>] | E<sub>max</sub> (%DA) | Log [EC<sub>50</sub>] |
| 0 M | 100.00 ± 1.75 | -6.88 ± 0.05 | 100.00 ± 2.98 | -5.42 ± 0.06 |
| 100.0 pM | 98.49 ± 1.49 | -6.89 ± 0.04 | 93.60 ± 2.39 | -5.47 ± 0.05 |
| 1.0 nM | 99.05 ± 1.56 | -6.91 ± 0.05 | 88.85 ± 2.23* | -5.67 ± 0.05* |
| 10.0 nM | 102.40 ± 1.60 | -7.07 ± 0.05 | 87.63 ± 2.10* | -5.89 ± 0.05* |
| 100.0 nM | 105.80 ± 2.02 | -7.29 ± 0.07 | 102.10 ± 2.09 | -6.19 ± 0.05* |
| 1.0 μM | 105.60 ± 1.98 | -7.56 ± 0.08* | 105.80 ± 2.38 | -6.39 ± 0.06* |
| 10.0 μM | 107.50 ± 2.00* | -7.72 ± 0.07* | 134.60 ± 1.89* | -6.66 ± 0.04* |
| 100.0 μM | 108.40 ± 1.80* | -7.79 ± 0.06* | 146.10 ± 1.86* | -7.06 ± 0.04* |

Data were presented as means ± SEM (n ≥ 3). \*p<0.05 compared to DA alone within each column using one-way ANOVA followed by post hoc Dunnett test.

**Supplementary Table 2.** Bias factor calculation between Gα_s_ and β-arrestin2.

| DA + [LY31] | Δlog | $\Delta\Delta$log |
| --- | --- | --- |
| 0 M | 1.46 |  |
| 100.0 pM | 1.44 | -0.02 |
| 1.0 nM | 1.29 | -0.17 |
| 10.0 nM | 1.25 | -0.21 |
| 100.0 nM | 1.12 | -0.34 |
| 1.0 μM | 1.17 | -0.29 |
| 10.0 μM | 0.96 | -0.50 |
| 100.0 μM | 0.60 | -0.86 |

**Supplementary Table 3.** LY3154207 differentially potentiated DA-induced G-protein coupling in D1R.

| DA + [LY31] | G$\alpha_s$ engagement | | G$\alpha_{olf}$ engagement | |
| --- | --- | --- | --- | --- |
|  | E<sub>max</sub> (%DA) | Log [EC<sub>50</sub>] | E<sub>max</sub> (%DA) | Log [EC<sub>50</sub>] |
| 0 M | 100.00 ± 8.09 | -5.17 ± 0.15 | 100.00 ± 7.00 | -5.60 ± 0.16 |
| 100.0 pM | 101.40 ± 9.48 | -5.10 ± 0.16 | 86.56 ± 7.60 | -5.62 ± 0.18 |
| 1.0 nM | 101.80 ± 10.65 | -5.07 ± 0.17 | 84.11 ± 6.28 | -5.64 ± 0.14 |
| 10.0 nM | 94.77 ± 10.39 | -5.17 ± 0.20 | 86.26 ± 6.77 | -5.76 ± 0.15 |
| **100.0 nM** | 120.80 ± 6.14 | -5.93 ± 0.12* | 100.90 ± 5.71 | -6.30 ± 0.15* |
| **1.0 μM** | 117.30 ± 6.50 | -5.97 ± 0.13* | 94.30 ± 4.25 | -6.54 ± 0.12* |

Data were presented as means ± SEM (n ≥ 3). \*p<0.05 compared to DA alone within each column using one-way ANOVA followed by post hoc Dunnett test.

**Supplementary Table 4.** LY3154207 differentially potentiated DA-induced cAMP production mediated by Gα_s_ and Gα _olf_.

|  | **D1R-$G\alpha_s$** |  | **D1R-$G\alpha_{olf}$** |  |
| --- | --- | --- | --- | --- |
| **DA + [PAM]** | $E_{max}$ (%DA) | Log [EC<sub>50</sub>] | $E_{max}$ (%DA) | Log [EC<sub>50</sub>] |
| **0 M** | 100.00 ± 3.17 | -8.44 ± 0.11 | 100.00 ± 3.55 | -8.35 ± 0.12 |
| **0.1 nM** | 110.20 ± 3.40 | -8.50 ± 0.11 | 102.50 ± 4.11 | -8.36 ± 0.14 |
| **1.0 nM** | 109.60 ± 3.50 | -8.56 ± 0.12 | 102.80 ± 3.96 | -8.33 ± 0.13 |
| **10.0 nM** | 113.50 ± 2.85 * | -8.64 ± 0.10 | 107.90 ± 4.23 | -8.35 ± 0.14 |
| **100.0 nM** | 118.00 ± 2.61 ** | -8.94 ± 0.11 | 108.10 ± 4.05 | -8.76 ± 0.16 |
| **1.0 μM** | 113.40 ± 2.90 * | -8.99 ± 0.19 | 108.50 ± 3.19 | -8.90 ± 0.17 |

Data were presented as means ± SEM (n ≥ 3). \*p<0.05 compared to DA alone within each column using one-way ANOVA followed by post hoc Dunnett test.

**Supplementary Table 5.** LY3154207 differentially potentiated DHX-induced cAMP production mediated by Gα_s_ and Gα _olf_.

|  | **D1R-$G\alpha_s$** |  | **D1R-$G\alpha_{olf}$** |  |
| --- | --- | --- | --- | --- |
| **DHX + [PAM]** | $E_{max}$ (%DA) | Log [EC<sub>50</sub>] | $E_{max}$ (%DA) | Log [EC<sub>50</sub>] |
| **0 M** | 100.00 ± 2.25 | -9.61 ± 0.09 | 100.00 ± 3.74 | -9.48 ± 0.15 |
| **0.1 nM** | 99.89 ± 2.43 | -9.60 ± 0.10 | 101.30 ± 3.80 | -9.41 ± 0.15 |
| **1.0 nM** | 98.74 ± 2.73 | -9.63 ± 0.11 | 99.21 ± 3.86 | -9.66 ± 0.16 |
| **10.0 nM** | 99.19 ± 3.38 | -9.79 ± 0.17 | 97.86 ± 3.62 | -9.82 ± 0.17 |
| **100.0 nM** | 104.00 ± 2.73 | -10.00 ± 0.22 | 99.93 ± 3.43 | -10.15 ± 0.24 |
| **1.0 μM** | 105.30 ± 2.61 | -10.12 ± 0.24 | 99.19 ± 3.47 | -10.18 ± 0.30 |

Data were presented as means ± SEM (n ≥ 3). \*p<0.05 compared to DA alone within each column using one-way ANOVA followed by post hoc Dunnett test.

**Supplementary Table 6.**
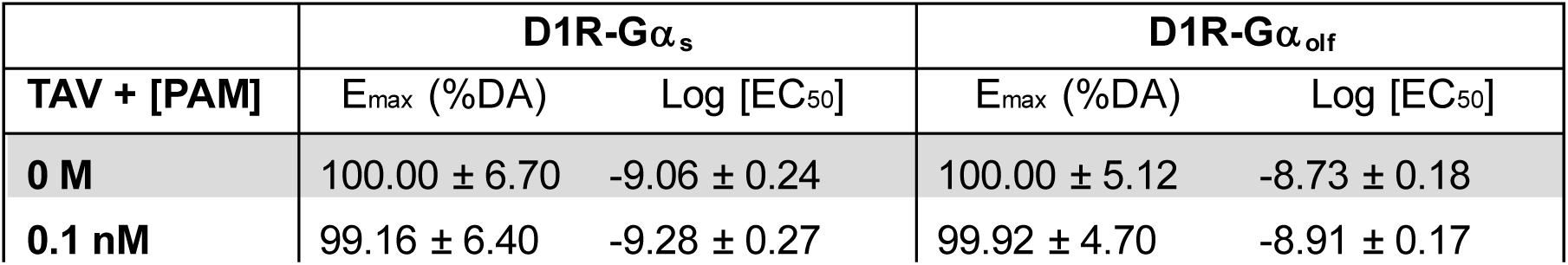
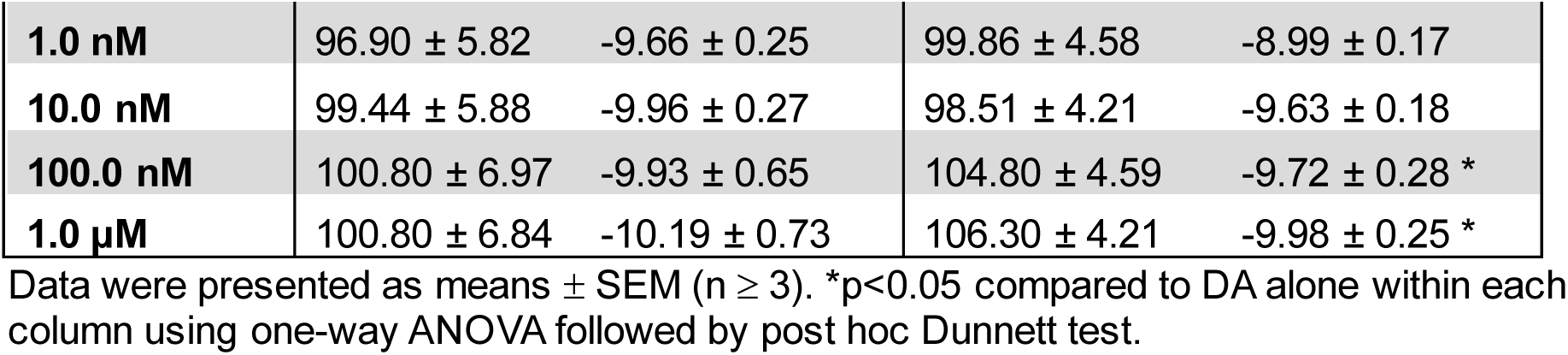
LY3154207 differentially potentiated TAV-induced cAMP production mediated by Gα_s_ and Gα _olf_.

**Supplementary Table 7.** W123 and R130 in ICL2 are crucial for LY3154207 to potentiate the DA-induced Gα_s_-mediated cAMP production by D1R.

|  | **D1R W123A-$G\alpha_s$** |  | **D1R R130A-$G\alpha_s$** |  |
| --- | --- | --- | --- | --- |
| **DA + [PAM]** | $E_{max}$ (%DA) | Log [EC<sub>50</sub>] | $E_{max}$ (%DA) | Log [EC<sub>50</sub>] |
| **0 nM** | 100.00 ± 2.90 | -8.39 ± 0.10 | 100.00 ± 2.16 | -8.60 ± 0.08 |
| **0.1 nM** | 103.40 ± 3.52 | -8.27 ± 0.11 | 100.40 ± 2.01 | -8.60 ± 0.07 |
| **1.0 nM** | 101.10 ± 3.33 | -8.35 ± 0.11 | 101.80 ± 1.69 | -8.63 ± 0.06 |
| **10.0 nM** | 100.00 ± 3.36 | -8.34 ± 0.12 | 94.82 ± 2.15 | -8.68 ± 0.08 |
| **100.0 nM** | 100.70 ± 3.42 | -8.40 ± 0.12 | 95.10 ± 2.24 | -8.62 ± 0.08 |
| **1.0 µM** | 103.80 ± 3.10 | -8.35 ± 0.11 | 98.68 ± 2.13 | -8.91 ± 0.10 |

Data were presented as means ± SEM (n ≥ 3). \*p<0.05 compared to DA alone within each column using one-way ANOVA followed by post hoc Dunnett test.

**Supplementary Table 8.** W123 and R130 in ICL2 are crucial for LY3154207 to potentiate the DA-induced Gα _olf_-mediated cAMP production by D1R.

|  | **D1R W123A-$G\alpha_{olf}$** |  | **D1R R130A-$G\alpha_{olf}$** |  |
| --- | --- | --- | --- | --- |
| **DA + [PAM]** | $E_{max}$ (%DA) | Log [EC<sub>50</sub>] | $E_{max}$ (%DA) | Log [EC<sub>50</sub>] |
| **0 nM** | 100.00 ± 1.70 | -8.76 ± 0.06 | 100.00 ± 2.66 | -8.82 ± 0.09 |
| **0.1 nM** | 99.74 ± 1.79 | -8.86 ± 0.06 | 99.39 ± 2.50 | -8.88 ± 0.09 |
| **1.0 nM** | 100.10 ± 1.82 | -8.89 ± 0.06 | 101.30 ± 2.76 | -8.72 ± 0.09 |
| **10.0 nM** | 100.70 ± 1.90 | -8.78 ± 0.07 | 101.50 ± 3.00 | -8.64 ± 0.10 |
| **100.0 nM** | 98.33 ± 1.90 | -8.77 ± 0.07 | 98.93 ± 2.78 | -9.02 ± 0.10 |
| **1.0 µM** | 96.45 ± 1.95 | -9.02 ± 0.07 | 98.34 ± 2.56 | -9.32 ± 0.13* |

Data were presented as means ± SEM (n ≥ 3). \*p<0.05 compared to DA alone within each column using one-way ANOVA followed by post hoc Dunnett test.

**Supplementary Table 9.** The αN domain of G-protein contributed to the subtype selectivity of LY3154207.

|  | **D1R-SOaN** |  | **D1R-OSaN** |  |
| --- | --- | --- | --- | --- |
| **DA + [PAM]** | $E_{max}$ (%DA) | Log [EC<sub>50</sub>] | $E_{max}$ (%DA) | Log [EC<sub>50</sub>] |
| **0 nM** | 100.00 ± 4.14 | -8.93 ± 0.15 | 100.00 ± 3.29 | -8.74 ± 0.12 |
| **0.1 nM** | 100.40 ± 4.20 | -8.98 ± 0.15 | 99.61 ± 3.37 | -8.82 ± 0.12 |
| **1.0 nM** | 97.84 ± 4.18 | -9.17 ± 0.15 | 99.39 ± 3.32 | -8.90 ± 0.11 |
| **10.0 nM** | 96.83 ± 4.03 | -9.21 ± 0.17 | 96.20 ± 3.45 | -9.07 ± 0.13 |
| **100.0 nM** | 100.10 ± 3.46 | -9.35 ± 0.21 | 97.80 ± 3.68 | -9.36 ± 0.18 |
| **1.0 µM** | 98.82 ± 3.36 | -9.68 ± 0.24 | 96.12 ± 3.94 | -9.45 ± 0.23* |

Data were presented as means ± SEM (n ≥ 3). \*p<0.05 compared to DA alone within each column using one-way ANOVA followed by post hoc Dunnett test.

